# Emotion Down- and Up-Regulation Act on Spatially Distinct Brain Areas: Interoceptive Regions to Calm Down and Other Affective Regions to Amp Up

**DOI:** 10.1101/2021.09.20.461138

**Authors:** Jungwon Min, Kaoru Nashiro, Hyun Joo Yoo, Christine Cho, Padideh Nasseri, Shelby L. Bachman, Shai Porat, Julian F. Thayer, Catie Chang, Tae-ho Lee, Mara Mather

**Affiliations:** University of Southern California (CA 90089); University of California, Irvine (CA 92697); Vanderbilt University (TN 37235); Virginia Polytechnic Institute and State University (VA 24061)

## Abstract

Prior studies on emotion regulation identified a set of brain regions specialized for generating and controlling affect. Researchers generally agree that when up- and down-regulating emotion, control regions in the prefrontal cortex turn up or down activity in affect-generating areas. However, the assumption that turning up and down emotions produces opposite effects in the same affect-generating regions is untested. We call this assumption the ‘affective dial hypothesis.’ Our study tested this hypothesis by examining the overlap between the sets of regions activated during up-regulation and those deactivated during down-regulation in a large number of participants (N=105). We found that up- and down-regulation both recruit regulatory regions such as the inferior frontal gyrus and dorsal anterior cingulate gyrus but act on distinct affect-generating regions. While up-regulation increases BOLD signal in regions associated with emotion such as the amygdala, anterior insula, striatum and anterior cingulate gyrus as well as in regions associated with sympathetic vascular activity such as periventricular white matter, down-regulation decreases signal in regions receiving interoceptive input such as the posterior insula and postcentral gyrus. These findings indicate that up- and down-regulation do not generally exert opposing effects on the same affect-generating regions. Instead, they target different brain circuits.

**Significance Statement:** Many contexts require modulating one’s own emotions. Identifying the brain areas implementing these regulatory processes should advance understanding emotional disorders and designing potential interventions. The emotion regulation field has an implicit assumption we call the affective dial hypothesis: that both emotion up- and down-regulation modulate the same emotion-generating brain areas. Countering the hypothesis, our findings indicate that up- and down-modulating emotions target different brain areas. Thus, the mechanisms underlying emotion regulation differ more than previously appreciated for up- versus down-regulation. In addition to their theoretical importance, these findings are critical for researchers attempting to target activity in particular brain regions during an emotion regulation intervention.

## Introduction

As humans, we are able to strategically modulate our own emotions. Often, this involves diminishing negative emotions and intensifying positive emotions. But there are also situations when one would want to increase the intensity of negative emotions (such as when wanting to feel empathy for a friend’s grief) or decrease the intensity of positive emotions (such as when trying not to laugh at a child’s embarrassing mistake). Thus, both diminishing and intensifying are processes that operate across valence and type of emotions (Gross, 2015).

Prior neuroimaging research indicates that diminishing and intensifying emotion rely on a shared set of affect-controlling regions that modulate activity in affect-generating regions (Buhle et al., 2014; Ochsner, Silvers, & Buhle, 2012). This set of control regions includes the ventrolateral, dorsomedial and dorsolateral prefrontal cortices (vlPFC, dmPFC, dlPFC), that are jointly recruited by up- and down-regulation and thus constitute an affect-control system (Kohn et al., 2014; Morawetz, Bode, Derntl, & Heekeren, 2017; Ochsner et al., 2012). On the other hand, the amygdala, insula, and striatum have been identified as affect-generating regions (Craig, 2009; Grosse Rueschkamp, Brose, Villringer, & Gaebler, 2019; Phelps, 2006), which can be up- or down-modulated by the control system (Braunstein, Gross, & Ochsner, 2017; Ochsner et al., 2012).

Despite its wide acceptance, the idea of the control system’s dialing up or down activity in affect-generating regions relies on an untested assumption: up-regulating (i.e., trying to intensify one’s emotions) will increase activity in the same affect-generating brain regions that down-regulating (i.e., trying to diminish one’s emotions) will decrease activity in. We call this implicit assumption of the emotion regulation field the **affective dial hypothesis** (see Figure 1).

**Figure 1.**
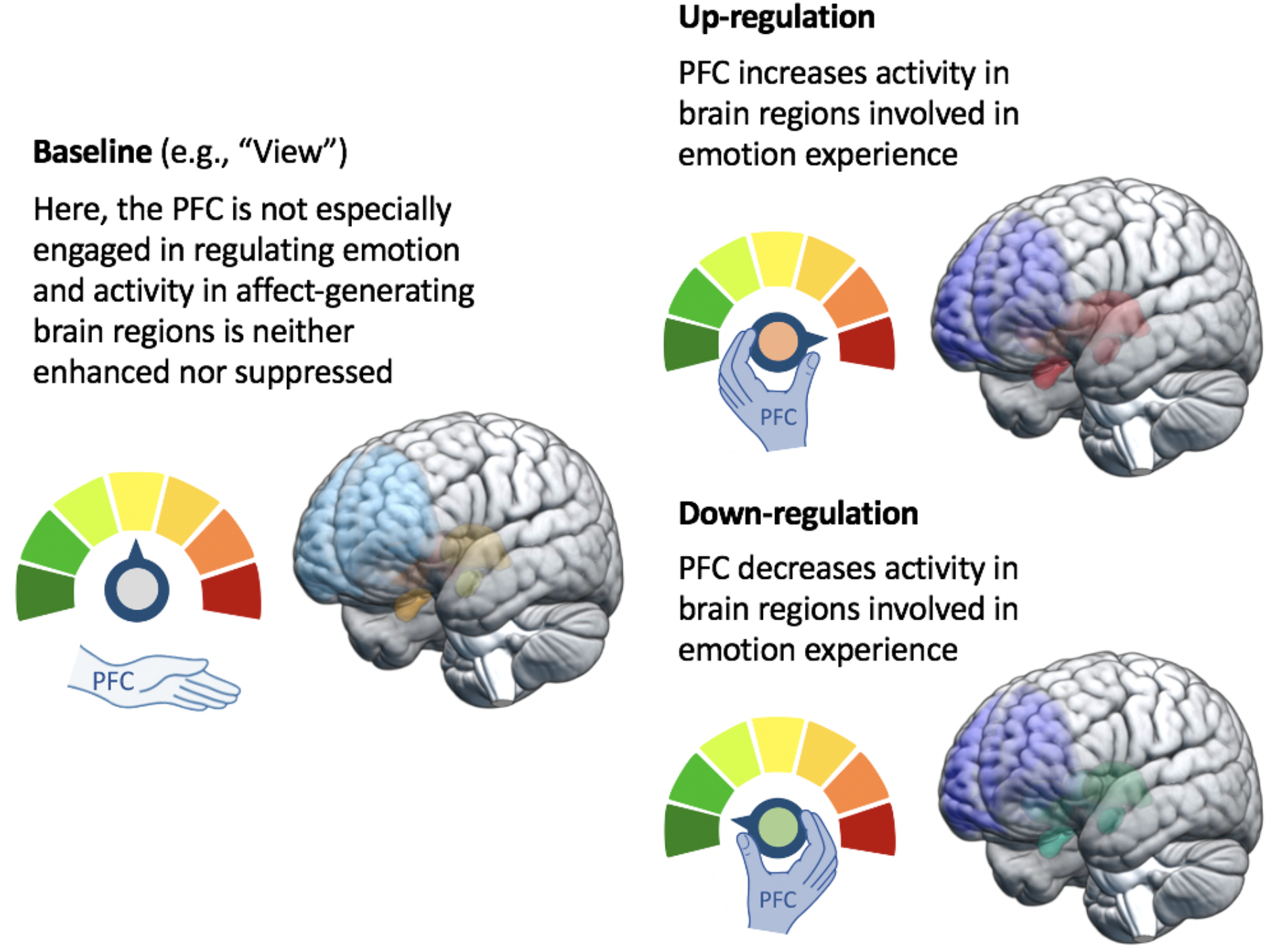
Schematic View of the Affective Dial Hypothesis *Note.* The control system (hand) dials down activity in affect-generating brain regions during emotion down-regulation and dials up activity in these same target regions during up-regulation. Simply viewing emotional images activates affect-generating brain regions without the action of the control system.

We found ten studies on young adults that included both up- and down-regulation trials as well as a non-regulation control (Domes et al., 2010; Eippert et al., 2007; Kim & Hamann, 2007; Leiberg, Eippert, Veit, & Anders, 2012; Li et al., 2018; Morawetz, Alexandrowicz, & Heekeren, 2017; Morawetz, Bode, Baudewig, Jacobs, & Heekeren, 2016; Morawetz, Bode, Baudewig, Kirilina, & Heekeren, 2016; Ochsner et al., 2004; Steinfurth et al., 2018), most of which were reported in a recent meta-analysis (Morawetz, Bode, et al., 2017). These studies typically showed increased activity in the vlPFC, dlPFC, supplementary motor area and anterior cingulate cortex during both intensifying and diminishing emotion. Furthermore, five of these studies conducted an explicit test of which regions were involved in regulation in both conditions by examining where overlap occurred between the up-regulation > baseline and down-regulation > baseline contrasts. All five of these studies showed some overlap between these two contrasts. Thus, this overlapping set of regions are involved in emotional control regardless of whether people are trying to up- or down-regulate their emotions. However, the affective dial idea that the same affect-generating regions are targeted by up- and down-regulation currently lacks support. None of those ten studies reported an explicit test of the overlap between up-regulate > baseline and baseline > down-regulate contrasts. Although six of the studies reported the baseline > down-regulate contrast at a whole-brain level, there were no consistently activated clusters. Thus, we could not find clear evidence that supports the affective dial hypothesis.

The current study tests the affective dial hypothesis that up-regulating emotions increases activity in the same affect-generating brain regions that down-regulating emotions decreases activity in. By having both up- and down-regulation types in one study and contrasting them with viewing trials, we directly tested how much the targets of up- and down-regulation overlap. In addition to whole-brain analyses, we included a region-of-interest (ROI) analysis of the amygdala, as many studies suggest it is a target of prefrontal regulatory systems (e.g., Berboth & Morawetz, 2021). We also investigated how the brain bases of the subjective sense of emotional experience differed during up- vs. down-regulation. Our sample size (N=105) gave us greater statistical power than prior studies comparing up- and down-regulating conditions.

## Methods

### Participants

The emotion regulation task was conducted as part of a 5-week heart rate variability biofeedback intervention study in which participants learned to modulate their heart rate by breathing at a slow rate (ClinicalTrials.gov Identifier: NCT03458910). The emotion regulation task was conducted both before and after the intervention, but for this paper we just used the baseline data from young adults before any intervention was conducted. Participants were recruited via USC’s subject pool, USC’s online bulletin board, Facebook, and flyers, and screened out for medical or psychiatric illnesses. However, people taking antidepressant or antianxiety medication were excluded only if they anticipated a change in treatment during the intervention. Upon completion or termination, participants were monetarily rewarded based on the total participation time and performance. As the present analyses focused on the pre-intervention session, we included participants who dropped out after the first emotion regulation task. This yielded 105 participants who ranged in age from 18 to 31 years (M_age_ = 22.8, SD_age_ = 2.69) and consisted of 54 males and 51 females.

### Task

We based our study design on a previously validated emotion regulation task (Kim & Hamann, 2007) which has up- and down-regulation trials for positive and negative emotions. We employed an event-related design. The 10-minute emotion regulation task had 42 trials, each of which consisted of a sequence involving a 1-second instruction, a 6-second regulation, and a 4-second rating period. During the 6-second regulation period, participants were asked to regulate emotion induced by the images according to the presented instruction. The instructions were “intensify,” “diminish,” or “view,” and the presented images were positive, negative, or neutral. Pairing of the instructions and images yielded 7 conditions: diminish-negative, diminish-positive, intensify-negative, intensify-positive, view-negative, view-positive, and view-neutral. After regulation, participants were asked to rate their strength of feeling with a scale from 1 (*weak*) to 4 (*strong*). Three trials from each condition were nested in a mini-block where the trials were separated by a fixation cross with a jittered interval that ranged from 0 to 4 seconds. The jittered intervals summed up to 4 seconds to keep the mini-block length the same, and the mini-blocks were spaced apart by a 5-second-long fixation cross. A total of 14 mini-blocks were arranged in a pseudorandom manner such that no blocks with the same instruction or image valence were shown consecutively. Six sets of images were selected from the International Affective Picture System such that the 18 negative, 18 positive, and 6 neutral images within each image set each had the same average valence (*M*_negative_ = 2.3, *M*_positive_ = 7.2, *M*_neutral_ = 5.0) and arousal scores (*M*_negative_ = 5.4, *M*_positive_ = 5.4, *M*_neutral_ = 2.8). During the task, each participant was presented with one of the six sets of images in a randomized order.

### Procedure

Participants had a practice session where they came up with their own reappraisal strategies to amplify, moderate, or passively experience the image-induced emotion according to the “intensify,” “diminish,” or “view” instruction. If they had difficulty devising their own method, they were presented with examples such as reinterpreting the situations or changing the distance between themselves and the scene. We also advised them not to generate an emotion opposite to the one that they were experiencing. For example, they were not supposed to replace a negative feeling with a positive one to diminish negative emotion. After the scan, participants were asked to report what regulation strategies they used and how successful they were in regulating emotions. For the four emotion-regulating conditions (e.g., diminish positive), 96% – 99% of participants used cognitive reappraisal and 92% – 98% of participants reported medium or high levels of confidence in their emotion regulation success.

### MRI data acquisition

MRI scans were conducted at USC’s Dana and David Dornsife Cognitive Neuroimaging Center using a 3T Siemens MAGNETOM Prisma MRI scanner with a 32-channel head coil. We obtained a T1-weighted MPRAGE anatomical image (TR = 2,300 ms, TE = 2.26 ms, slice thickness = 1.0 mm, flip angle = 9°, field of view = 256 mm, voxel size = 1.0 mm isotropic). We acquired 250 whole brain volumes of T2*-weighted functional images using multi-echo planar imaging sequence (TR= 2,400 mm, TE 18/35/53 ms, slice thickness = 3.0 mm, flip angle = 75°, field of view = 240 mm, voxel size = 3.0 mm isotropic).

### MRI data analysis

One hundred and fourteen participants completed the emotion regulation task during the baseline session. We excluded three participants whose multi-echo denoising process failed and six participants who failed to respond to more than 50% of the trials. This left 105 participants for fMRI analyses.

The functional MRI data were denoised with multi-echo independent component analysis which removed artifact components using the linear echo-time dependence of blood oxygen level dependent (BOLD) signal changes (Kundu, Inati, Evans, Luh, & Bandettini, 2012). The denoised data was entered into FMRIB Software Library (FSL) version 6.0 for the individual- and group-level analysis. Individual-level analysis included two steps of affine linear transformation with 12 degrees of freedom where each functional image was registered to the MNI152 T1 2mm template via its T1-weighted anatomical image. Individual-level analysis also included a preprocessing of motion correction, spatial smoothing with 5 mm FWHM, and high-pass filtering with 600-second cutoff. Individual whole-brain BOLD time series were modelled with a linear combination of seven emotion-regulation regressors during the 6-second emotion regulation period (diminish-negative, view-negative, intensify-negative, diminish-positive, view-positive, intensify-positive, and view-neutral) along with their temporal derivatives, each convolved with a double-gamma hemodynamic response function. For the group-level analysis, FSL’s mixed-effects model (FLAME 1) was used to test the mean effect of emotion regulation, contrasted across the conditions. The final results were corrected for family-wise error at *p* < .05 with the cluster-wise threshold at *z* > 3.1. We tested for overlapping control regions via a conjunction analysis taking the intersection of intensify > view and diminish > view and tested the affective dial hypothesis via a conjunction analysis taking the intersection of intensify > view and view > diminish.

To characterize the nature of the brain areas identified by the view > diminish and intensify > view contrasts, we used emotion-associated and interoception-associated cluster maps from a prior meta-analytic study (Adolfi et al., 2017). We derived three maps from this meta-analysis study: 1) the intersection of the two meta-analytic maps; 2) the emotion-associated map with the intersection regions removed; and 3) the interoception-associated map with the intersection regions removed. We then overlapped these three meta-analytic maps with the thresholded view > diminish and intensify > view contrast maps (after removing the intersection of diminish > view and intensify > view to remove activity likely related to regulation effort rather than its effects), counted the number of voxels overlapping each of the three meta-analytic maps, and divided the number of overlapping voxels with the total number of voxels in each thresholded contrast map.

To assess the BOLD activity changes in the amygdala, we individually segmented the amygdala region from each participant’s T1-weighted image using FreeSurfer version 6 and created the left and right amygdala masks in the native space. We then applied FSL FLIRT to transform the masks to the standard MNI space and input them to Featquery to obtain average percent signal change values in the amygdala activity during emotion regulation.

Subjective ratings during the task were analyzed by using SPSS to conduct an ANOVA with mean emotional intensity as the dependent variable and the regulation goals (diminish, intensify, view) and image valence (negative, positive) as within-subject independent factors.

We also examined how brain activity while implementing different regulation goals relates to subjective ratings of emotion regulation outcome. To do so, we normalized the online rating scores within each subject’s intensify or diminish condition and used the normalized scores as a weight for the two emotion-regulation regressors (diminish, intensify; each aggregated across positive and negative valence) in another individual-level analysis. We excluded nine subjects who always responded with the same rating within either condition, which made normalization impossible within that condition for that person. The subsequent group-level analysis tested the mean effect of four contrasts: diminish, intensify, diminish > intensify, and intensify > diminish.

## Results

### Subjective Ratings

There was a significant main effect of the three emotion regulation goals, *F*(2, 208) = 228.60, *r* = 0.83, *p* < 0.001 and of the emotional valence, *F*(1,104) = 5.58, *r* = 0.23, *p* = 0.02 on self-rated emotional intensity. But there was no significant interaction between goals and valence, *F*(2, 208) = 1.73, *r* = 0.13, *p* = 0.18. We also conducted Bonferroni-corrected t-tests for pairs of regulation and valence types. The corrected *p* threshold was at 0.007. Subjective intensity ratings were higher for intensifying than for viewing, *t*(104) = 12.68, *r* = 0.61, *p* < 0.001 for negative emotion and *t*(104) = 16.19, *r* = 0.63, *p* < 0.001 for positive emotion, and also higher for viewing than for diminishing, *t*(104) = 5.44, *r* = 0.29, *p* < 0.001 for negative emotion and *t*(104) = 5.09, *r* = 0.25, *p* < 0.001 for positive emotion (Figure 3). Ratings did not significantly differ between negative and positive emotion for either intensifying, *t*(104) = 0.60, *r* = 0.03, *p* = 0.55, for diminishing, *t*(104) = 2.43, *r* = 0.09, *p* = 0.02, or for viewing, *t*(104) = 2.46, *r* = 0.10, *p* = 0.02 though the comparisons were significant at an uncorrected level for diminishing and viewing (see Table 1 for details).

**Table 1.** Subjective Ratings across Regulation and Valence

|  | Negative |  |  | Positive |  |  |
| --- | --- | --- | --- | --- | --- | --- |
|  | M | SE | 95% CI | M | SE | 95% CI |
| Diminish | 1.95 | 0.06 | [1.82, 2.07] | 1.83 | 0.07 | [1.7, 1.96] |
| View | 2.33 | 0.07 | [2.2, 2.46] | 2.19 | 0.07 | [2.05, 2.32] |
| Intensify | 3.27 | 0.05 | [3.16, 3.37] | 3.23 | 0.06 | [3.12, 3.35] |

**Figure 2.**
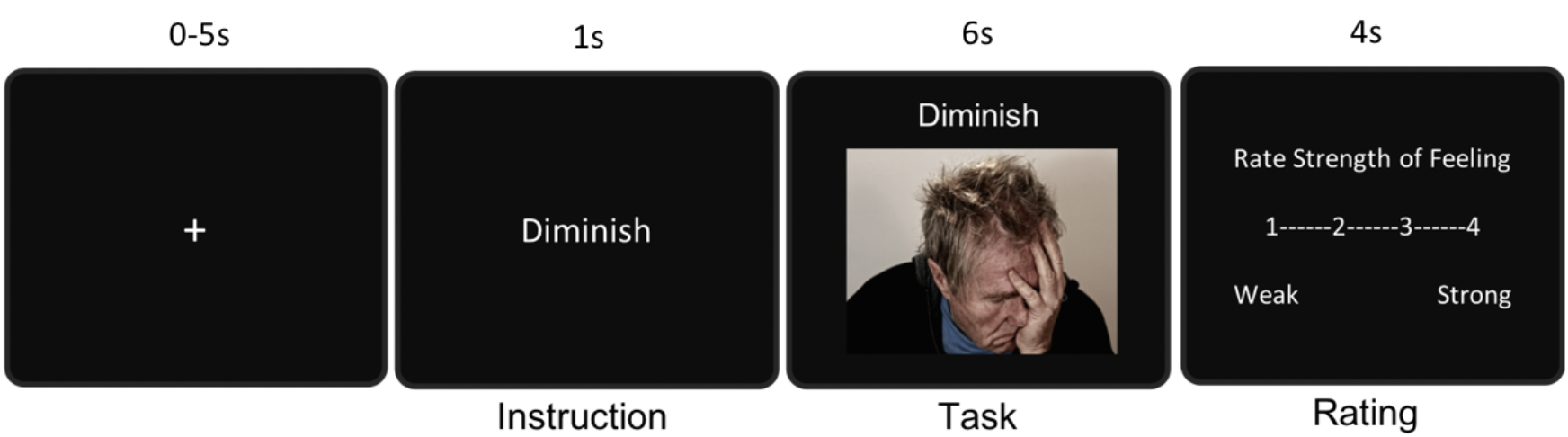
Emotion Regulation Trial Design

**Figure 3.**
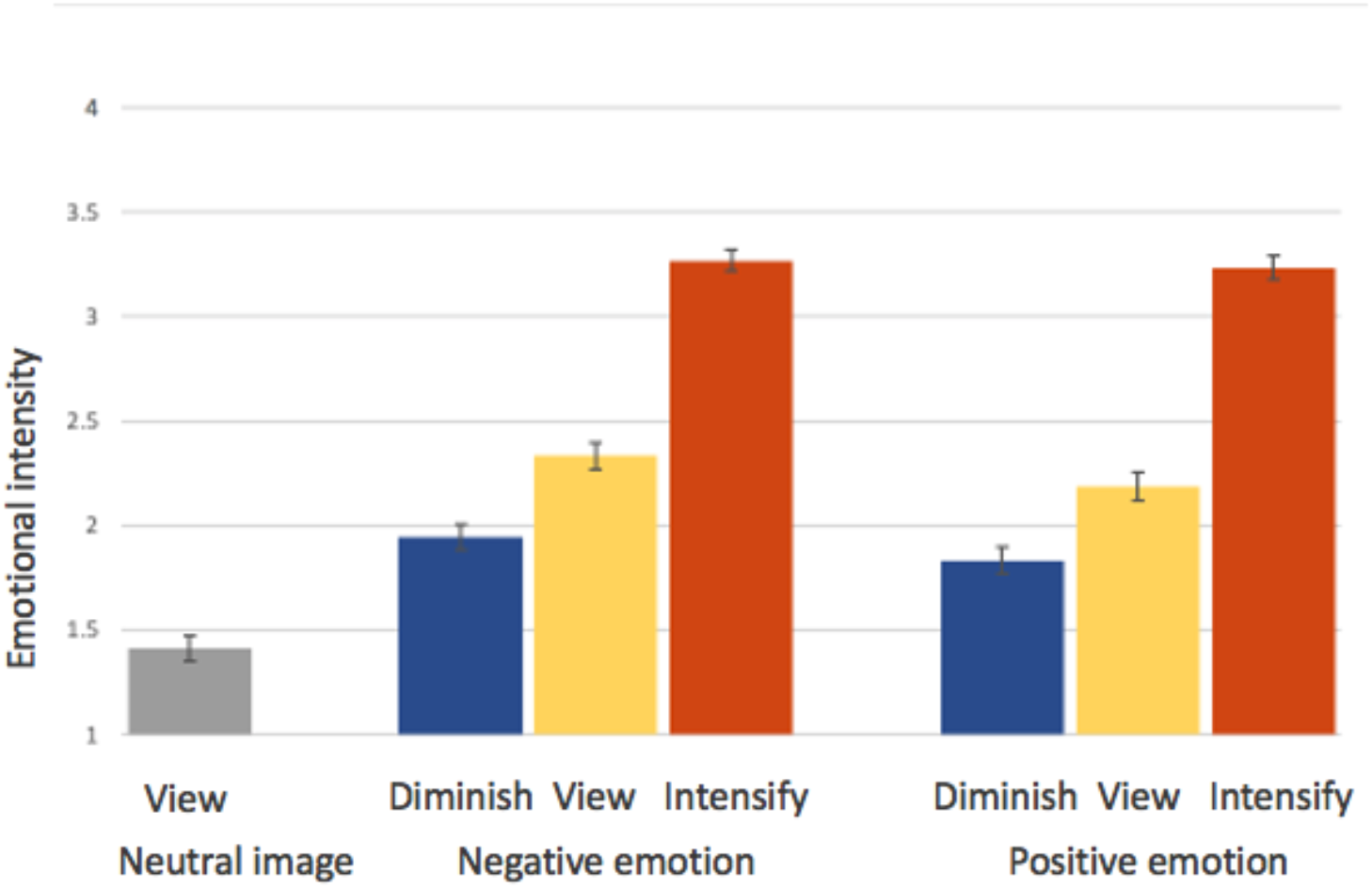
Subjective Ratings of Emotional Intensity *Note.* The error bars reflect the standard error of each condition.

### Regulation Effort

Our analyses focused on the general regulatory effect of emotion regulation across positive and negative valence, based on prior findings that the brain’s affective workspace varies little across valence (Lindquist, Satpute, Wager, Weber, & Barrett, 2016). Contrasting the diminish against view condition (diminish > view) revealed brain regions showing increased activation during emotional down-regulation (Figure 4A, Table 2): the anterior insular cortex, lateral frontal orbital cortex, dorsal anterior cingulate gyrus, paracingulate gyrus, superior frontal gyrus, and inferior frontal gyrus. Contrasting the intensify against view condition (intensify > view) revealed brain regions showing increased activation during emotional up-regulation (Figure 4B, Table 3): the anterior insular cortex, lateral frontal orbital cortex, frontal medial cortex, anterior cingulate gyrus, posterior cingulate gyrus, inferior frontal gyrus, middle frontal gyrus, superior frontal gyrus, hippocampus, amygdala, putamen, and thalamus. Consistent with prior studies (Domes et al., 2010; Eippert et al., 2007; Kim & Hamann, 2007; Li et al., 2018; Ochsner et al., 2004), there were a number of brain regions activated during both up- and down-regulation (intensify > view ∩ diminish > view), consistent with regulatory regions shared by the two opposing regulation goals. These regions were the insular cortex, inferior frontal gyrus, middle frontal gyrus, superior frontal gyrus, dorsal anterior cingulate gyrus, and angular gyrus (Figure 5A).

**Table 2.** List of Regions (Figure 4A) which Increased Activity during Down-regulation (diminish > view)

| Diminish > View Clusters<br>(Harvard-Oxford Structural Atlas) | MNI coordinate |  |  | Zmax | Voxels |
| --- | --- | --- | --- | --- | --- |
|  | x | y | z |  |  |
| Supplementary Motor Cortex | -2 | 6 | 60 | 5.96 | 221 |
| Paracingulate Gyrus | -2 | 14 | 50 | 5.36 | 202 |
| Angular Gyrus | -58 | -54 | 20 | 5.24 | 153 |
| Superior Frontal Gyrus | -4 | 12 | 58 | 5.23 | 230 |
| Frontal Pole | -26 | 50 | 32 | 4.99 | 445 |
| Cerebellum Right Crus I | 36 | -64 | -38 | 4.99 | 212 |
| Middle Frontal Gyrus | -46 | 14 | 38 | 4.89 | 291 |
| Frontal Operculum Cortex | -44 | 18 | 0 | 4.88 | 70 |
| Frontal Orbital Cortex | -38 | 20 | -14 | 4.74 | 335 |
| Inferior Frontal Gyrus, pars opercularis | -50 | 12 | 4 | 4.68 | 116 |
| Lateral Occipital Cortex, superior division | -48 | -64 | 40 | 4.63 | 194 |
| Cerebellum Right Crus II | 20 | -72 | -38 | 4.49 | 181 |
| Supramarginal Gyrus, posterior division | -60 | -48 | 24 | 4.46 | 86 |
| Cingulate Gyrus, anterior division | -4 | 20 | 34 | 4.17 | 52 |
| Inferior Frontal Gyrus, pars triangularis | -54 | 24 | 8 | 4.15 | 14 |
| Insular Cortex | -40 | 16 | -2 | 3.91 | 34 |
| Precentral Gyrus | -44 | -4 | 54 | 3.83 | 13 |
| Cerebellum Right VI | 10 | -76 | -18 | 3.76 | 23 |
| Lateral Occipital Cortex, inferior division | -54 | -66 | 12 | 3.64 | 5 |
| Lingual Gyrus | 12 | -76 | -10 | 3.55 | 20 |
| Cerebellum Right VIIb | 18 | -70 | -42 | 3.37 | 3 |
| Temporal Pole | -44 | 18 | -18 | 3.32 | 3 |

**Table 3.** List of Regions (Figure 4B) which Increased Activity during Up-regulation (intensify > view)

| Intensify > View Clusters<br>(Harvard-Oxford Structural Atlas) | MNI coordinate |  |  |  | Voxels |
| --- | --- | --- | --- | --- | --- |
|  | x | y | z | Zmax |  |
| Left Thalamus | -2 | -22 | 6 | 7.38 | 938 |
| Insular Cortex | -38 | 4 | 2 | 7.11 | 366 |
| Cingulate Gyrus, anterior division | -2 | 14 | 34 | 7.09 | 877 |
| Cerebellum Right Crus I | 30 | -76 | -36 | 6.68 | 1129 |
| Brain Stem | -2 | -32 | -4 | 6.66 | 537 |
| Left Hippocampus | -30 | -36 | -4 | 6.62 | 271 |
| Superior Frontal Gyrus | -12 | -2 | 70 | 6.43 | 109 |
| Central Opercular Cortex | -42 | 6 | 2 | 6.41 | 324 |
| Cerebellum Right Crus II | 30 | -76 | -38 | 6.39 | 875 |
| Frontal Operculum Cortex | -44 | 24 | 0 | 6.36 | 119 |
| Supplementary Motor Cortex | 4 | 0 | 68 | 6.28 | 498 |
| Right Thalamus | 2 | -8 | 6 | 6.28 | 468 |
| Precentral Gyrus | 54 | 0 | 44 | 6.20 | 297 |
| Temporal Pole | -48 | 18 | -16 | 6.18 | 538 |
| Lateral Occipital Cortex, superior division | -46 | -72 | 24 | 6.15 | 317 |
| Frontal Orbital Cortex | -44 | 24 | -6 | 6.04 | 184 |
| Supramarginal Gyrus, posterior division | -58 | -46 | 22 | 6.00 | 76 |
| Left Caudate | -16 | -8 | 20 | 5.95 | 373 |
| Cerebellum Right V | 2 | -62 | -6 | 5.91 | 47 |
| Left Lateral Ventricle | -14 | 24 | 4 | 5.87 | 733 |
| Left Pallidum | -12 | 4 | -4 | 5.87 | 102 |
| Cerebellum Vermis VI | 0 | -70 | -18 | 5.81 | 216 |
| Frontal Pole | -30 | 44 | 24 | 5.81 | 1466 |
| Cerebellum Left I-IV | -6 | -50 | -6 | 5.80 | 189 |
| Right Lateral Ventricle | 10 | -4 | 18 | 5.73 | 560 |
| Cerebellum Left Crus I | -42 | -56 | -40 | 5.68 | 518 |
| Cerebellum Left V | 0 | -60 | -6 | 5.65 | 184 |
| Right Caudate | 18 | -6 | 24 | 5.64 | 194 |
| Left Putamen | -30 | 4 | 4 | 5.50 | 400 |
| Angular Gyrus | -54 | -54 | 18 | 5.43 | 96 |
| Right Hippocampus | 32 | -36 | -6 | 5.37 | 105 |
| Middle Temporal Gyrus, anterior division | -54 | -4 | -28 | 5.28 | 88 |
| Left Accumbens | -6 | 12 | -4 | 5.27 | 53 |
| Precuneus Cortex | -14 | -58 | 18 | 5.26 | 373 |
| Cingulate Gyrus, posterior division | -4 | -54 | 28 | 5.26 | 197 |
| Cerebellum Right I-IV | 2 | -46 | -6 | 5.21 | 49 |
| Lingual Gyrus | -10 | -52 | -4 | 5.17 | 213 |
| Parahippocampal Gyrus, posterior division | -18 | -26 | -20 | 5.16 | 28 |
| Cerebellum Right VI | 8 | -74 | -22 | 5.15 | 252 |
| Parietal Operculum Cortex | -34 | -30 | 20 | 5.01 | 130 |
| Middle Frontal Gyrus | -34 | 30 | 44 | 4.95 | 171 |
| Cerebellum Right VIIb | 18 | -72 | -46 | 4.92 | 83 |
| Inferior Frontal Gyrus, pars opercularis | -50 | 12 | 4 | 4.86 | 113 |
| Frontal Medial Cortex | -6 | 54 | -10 | 4.82 | 64 |
| Cerebellum Right VIIIb | 14 | -42 | -54 | 4.81 | 15 |
| Planum Polare | -54 | 2 | -2 | 4.81 | 31 |
| Cerebellum Left VI | -14 | -62 | -26 | 4.78 | 281 |
| Cerebellum Left Crus II | -42 | -56 | -44 | 4.73 | 100 |
| Right Putamen | 18 | 10 | -8 | 4.73 | 66 |
| Planum Temporale | -60 | -36 | 16 | 4.67 | 31 |
| Paracingulate Gyrus | -4 | 18 | 38 | 4.67 | 207 |
| Middle Temporal Gyrus, temporo-occipital part | -60 | -56 | 2 | 4.67 | 130 |
| Temporal Fusiform Cortex, posterior division | -40 | -34 | -20 | 4.66 | 72 |
| Lateral Occipital Cortex, inferior division | -54 | -64 | 10 | 4.62 | 81 |
| Left Amygdala | -14 | -6 | -16 | 4.61 | 84 |
| Subcallosal Cortex | -2 | 12 | -4 | 4.57 | 54 |
| Temporal Occipital Fusiform Cortex | 34 | -46 | -8 | 4.56 | 14 |
| Cerebellum Right IX | 6 | -50 | -52 | 4.52 | 29 |
| Cerebellum Right VIIIa | 36 | -52 | -52 | 4.51 | 43 |
| Occipital Pole | -8 | -96 | 2 | 4.37 | 59 |
| Cerebellum Vermis IX | 2 | -52 | -32 | 4.34 | 12 |
| Superior Temporal Gyrus, posterior division | 66 | -30 | 14 | 4.28 | 27 |
| Parahippocampal Gyrus, anterior division | -18 | -20 | -24 | 4.07 | 25 |
| Cerebellum Vermis X | 2 | -50 | -34 | 4.02 | 7 |
| Cerebellum Left IX | -12 | -46 | -52 | 3.99 | 13 |
| Middle Temporal Gyrus, posterior division | -64 | -42 | -10 | 3.96 | 11 |
| Intracalcarine Cortex | -6 | -68 | 12 | 3.87 | 13 |
| Cerebellum Left X | -22 | -40 | -44 | 3.85 | 14 |
| Right Amygdala | 16 | -8 | -18 | 3.80 | 26 |
| Inferior Frontal Gyrus, pars triangularis | -52 | 24 | -2 | 3.79 | 19 |
| Occipital Fusiform Gyrus | 28 | -72 | -8 | 3.77 | 11 |
| Cerebellum Vermis VIIIa | 2 | -72 | -42 | 3.72 | 7 |
| Right Accumbens | 10 | 10 | -8 | 3.54 | 17 |
| Inferior Temporal Gyrus, anterior division | -48 | -2 | -34 | 3.37 | 2 |
| Cerebellum Left VIIIa | -30 | -44 | -48 | 3.35 | 5 |
| Postcentral Gyrus | -22 | -38 | 62 | 3.31 | 4 |
| Cerebellum Vermis Crus II | 0 | -78 | -30 | 3.30 | 2 |
| Supramarginal Gyrus, anterior division | -64 | -38 | 28 | 3.29 | 3 |
| Cerebellum Left VIIIb | -24 | -40 | -50 | 3.28 | 11 |
| Superior Temporal Gyrus, anterior division | -58 | 2 | -6 | 3.23 | 1 |

**Figure 4.**
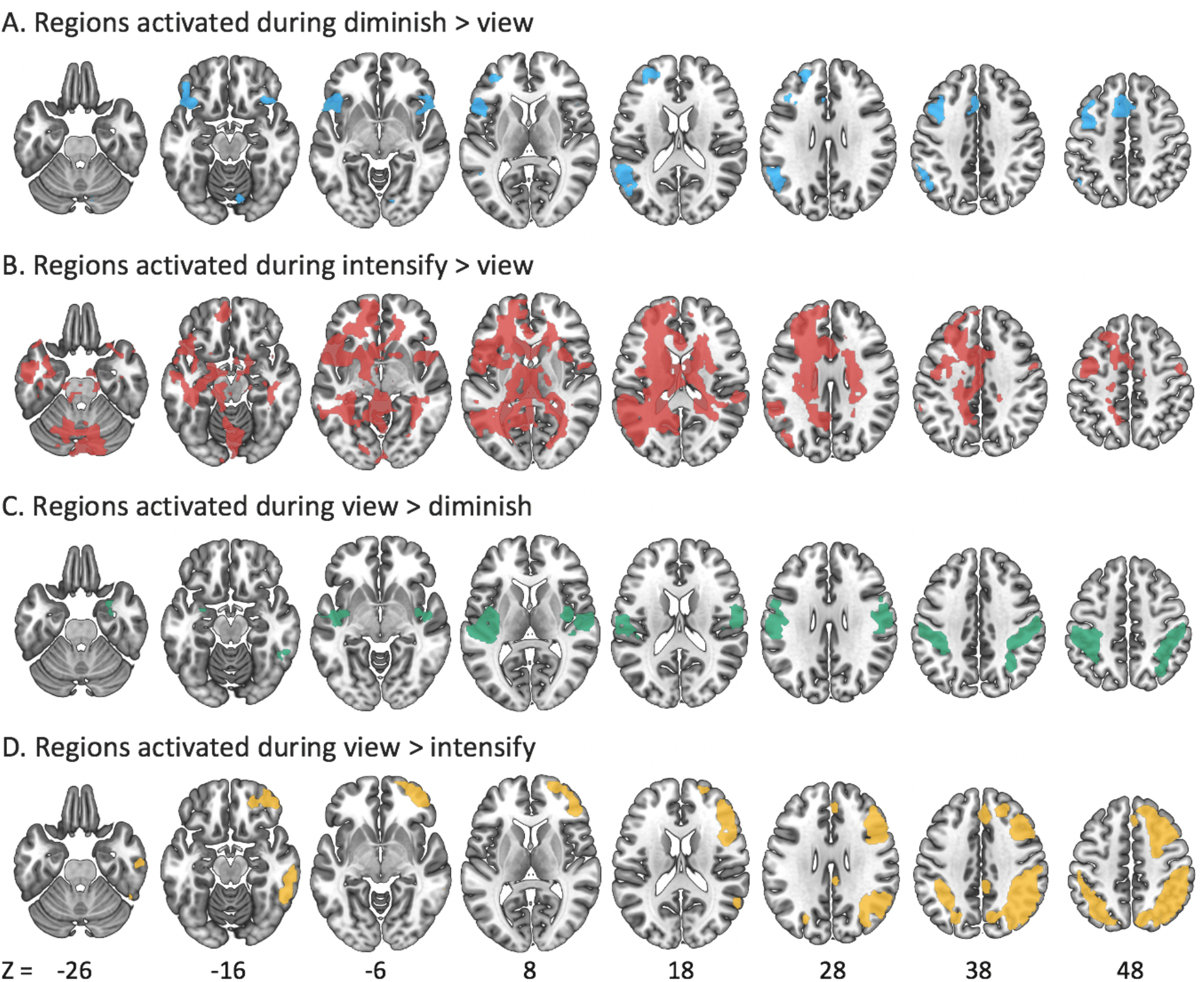
Regions Showing Activation Differences between View and Regulation Conditions. *Note.* (A) shows areas (blue) which increased activity during down-regulation, (B) shows areas (red) which increased activity during up-regulation, (C) shows areas (green) in which activity was decreased during down-regulation, and (D) shows areas (yellow) in which activity was decreased during up-regulation.

**Figure 5.**
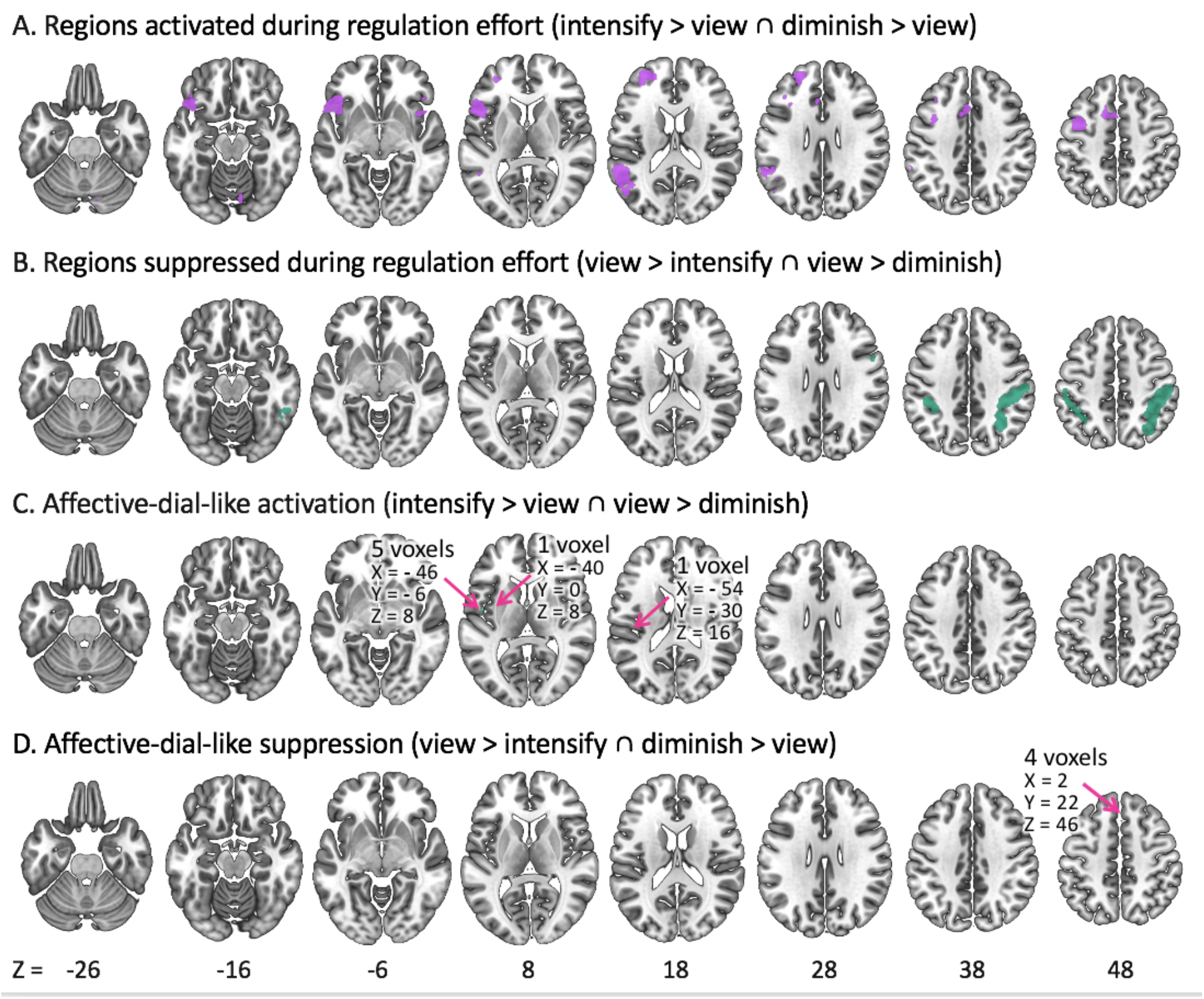
Brain Activity Consistent with Regulatory Effort vs. with Emotional Outcome Across Regulation Conditions *Note*. (A) shows common regions (purple) activated during both up- and down-regulation, while (B) shows regions (green) deactivated during both up- and down-regulation. (C) shows regions (turquoise) which increased activity during up-regulation and decreased activity during down-regulation, and (D) shows regions (orange) which decreased activity during up-regulation and increased activity during down-regulation.

To test the affective dial hypothesis, we examined the intersection of the two contrasts (intensify > view and view > diminish) that should show significant emotion-related activity if emotion regulation modulates affect-generating brain regions in the expected linear fashion (intensify > view > diminish). If emotion regulation processes act on the same affect-generating brain regions when up- and down-regulating, the intensify > view and view > diminish contrasts should show overlapping areas. Despite our robust power, however, there were only seven voxels that were significant for both the intensify > view and view > diminish contrasts. They were in the central opercular cortex (five voxels), the parietal operculum cortex (one voxel), and the insular cortex (one voxel). Besides these seven voxels (Figure 5C), there was no overlap between the significant clusters in the two contrasts, suggesting that up- and down-regulation act on two distinct emotion-generating networks. The intensify > view contrast (Figure 4B, Table 3) revealed the amygdala, striatum, anterior insular cortex and cingulate gyrus which are associated with emotional experience (Lindquist et al., 2016) as well as white matter and ventricular regions which are associated with vascular activity during sympathetic arousal (Özbay et al., 2019). The view > diminish contrast (Figure 4C, Table 4) showed the posterior insula cortex and postcentral gyrus, which receive visceral and sensory input and represent the physiological states of the body (Craig, 2002). The regions which lowered their activity during intensifying emotion (view > intensify) included the frontal pole, middle frontal gyrus, and angular gyrus (Figure 4D, Table 5). Similarly, examining the diminish > view and view > intensify intersection revealed only 4 voxels in the paracingulate gyrus consistent with a linear diminish > view > intensify affective-dial suppression pattern (Figure 5D).

**Table 4.** List of Regions (Figure 4C) which Decreased Activity during Down-regulation (view > diminish)

| View > Diminish Clusters<br>(Harvard-Oxford Structural Atlas) | MNI coordinate |  |  | Zmax | Voxels |
| --- | --- | --- | --- | --- | --- |
|  | x | y | z |  |  |
| Postcentral Gyrus | -42 | -32 | 60 | 5.75 | 761 |
| Superior Parietal Lobule | -36 | -50 | 60 | 5.92 | 363 |
| Insular Cortex | -40 | -6 | 8 | 6.38 | 352 |
| Central Opercular Cortex | -42 | -8 | 10 | 5.26 | 252 |
| Lateral Occipital Cortex, superior division | 32 | -64 | 44 | 4.46 | 236 |
| Supramarginal Gyrus, anterior division | -52 | -32 | 44 | 5.74 | 175 |
| Precentral Gyrus | -58 | 4 | 28 | 5.13 | 124 |
| Heschl's Gyrus including H1 and H2 | -46 | -24 | 12 | 5.08 | 111 |
| Inferior Temporal Gyrus, temporo-occipital part | 54 | -48 | -20 | 4.02 | 92 |
| Planum Temporale | -52 | -28 | 10 | 4.70 | 77 |
| Planum Polare | 48 | -8 | -6 | 4.69 | 57 |
| Parietal Operculum Cortex | -50 | -28 | 14 | 4.41 | 49 |
| Right Amygdala | 28 | 0 | -22 | 3.97 | 39 |
| Supramarginal Gyrus, posterior division | 50 | -38 | 54 | 4.88 | 18 |
| Left Amygdala | -28 | -4 | -16 | 3.63 | 16 |
| Temporal Pole | 28 | 6 | -28 | 3.38 | 10 |
| Parahippocampal Gyrus, anterior division | 22 | 4 | -32 | 3.38 | 5 |
| Right Hippocampus | 30 | -6 | -26 | 3.66 | 4 |
| Superior Temporal Gyrus, posterior division | 60 | -18 | -2 | 3.47 | 2 |
| Right Putamen | 32 | -10 | 6 | 3.44 | 2 |

**Table 5.** List of Regions (Figure 4D) which Decreased Activity during Up-regulation (view > intensify)

| View > Intensify Clusters<br>(Harvard-Oxford Structural Atlas) | MNI coordinate |  |  | Zmax | Voxels |
| --- | --- | --- | --- | --- | --- |
|  | x | y | z |  |  |
| Angular Gyrus | 48 | -56 | 48 | 7.77 | 420 |
| Cingulate Gyrus, posterior division | 4 | -36 | 32 | 5.17 | 148 |
| Frontal Orbital Cortex | 20 | 32 | -20 | 3.83 | 4 |
| Frontal Pole | 42 | 52 | -12 | 7.31 | 1725 |
| Inferior Frontal Gyrus, pars opercularis | 54 | 12 | 22 | 5.92 | 64 |
| Inferior Frontal Gyrus, pars triangularis | 54 | 30 | 16 | 4.02 | 3 |
| Inferior Temporal Gyrus, posterior division | 62 | -28 | -22 | 6.35 | 14 |
| Inferior Temporal Gyrus, temporooccipital part | 62 | -44 | -18 | 6.04 | 163 |
| Lateral Occipital Cortex, superior division | 40 | -60 | 46 | 9.24 | 1969 |
| Middle Frontal Gyrus | 36 | 16 | 52 | 7.18 | 439 |
| Middle Temporal Gyrus, posterior division | 66 | -24 | -18 | 6.00 | 94 |
| Middle Temporal Gyrus, temporooccipital part | 64 | -42 | -10 | 4.22 | 17 |
| Paracingulate Gyrus | 2 | 28 | 42 | 5.46 | 230 |
| Postcentral Gyrus | 54 | -22 | 46 | 5.21 | 104 |
| Precentral Gyrus | 54 | 10 | 24 | 6.39 | 50 |
| precuneus Cortex | 8 | -72 | 44 | 5.85 | 72 |
| Superior Frontal Gyrus | 24 | 24 | 56 | 6.33 | 157 |
| Superior Parietal Lobule | 42 | -46 | 56 | 6.03 | 119 |
| Supramarginal Gyrus, anterior division | 54 | -32 | 46 | 5.42 | 99 |
| Supramarginal Gyrus, posterior division | 50 | -44 | 50 | 7.15 | 119 |

The lack of much activity consistent with either an intensify > view > diminish or a diminish > view > intensify pattern suggests that intensifying and diminishing emotions target different brain networks to modulate emotion. To help characterize the nature of the brain regions which emotion up-versus down-regulation act on, we counted how many voxels activated during intensify > view versus view > diminish overlapped with emotion- versus interoception-associated cluster maps generated from a prior meta-analysis (Adolfi et al., 2017, see Figure 6 for maps). We found that 21.5% of activated voxels during view > diminish overlapped with interoception-related areas, while only 6.0% overlapped with emotion-related areas. During intensify > view, 15.9% overlapped emotion-related areas, while 5.7% overlapped interoception-related areas.

**Figure 6.**
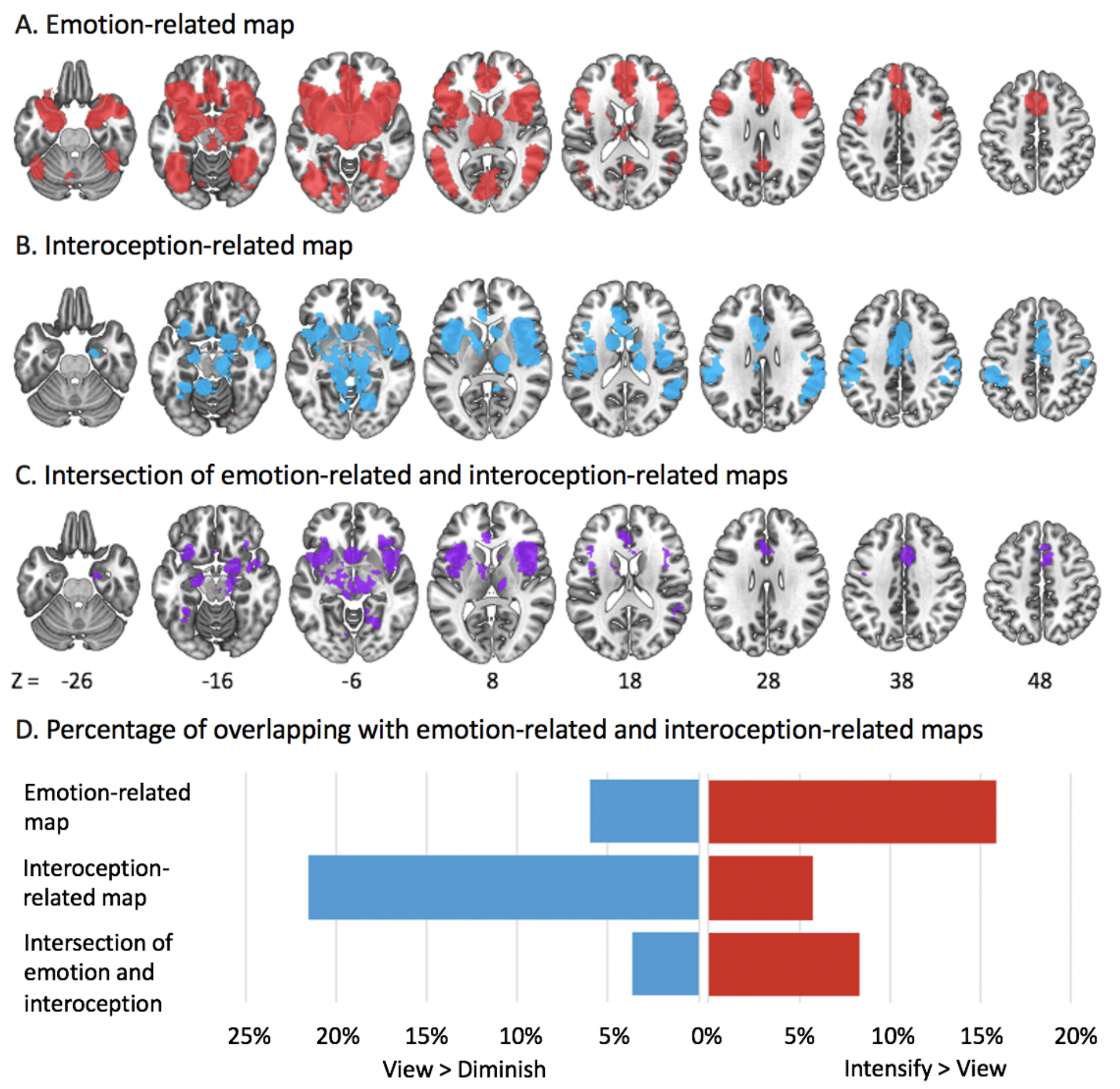
Emotion-related and interoception-related areas identified in Adolfi et al.’s meta-analysis *Note.* While clusters (red) in (A) are related to emotion, clusters (blue) in (B) are related to interoception. (C) is the intersection (purple) of (A) and (B). (D) shows the percentage of the voxels during down- and up-regulation (Figs. 4C, 4B) which overlap with (A), (B), and (C).

In our follow-up ROI analysis, although the amygdala numerically showed the affective-dial-like diminish < view < intensify pattern (Figure 7), neither the right or left amygdala showed both significant diminish < view and view < intensify effects as predicted by the affective dial hypothesis. A post-hoc t-test with Bonferroni-corrected *p* threshold at 0.01 showed that activity in the left amygdala differed between intensify and view, *t*(104) = 4.12, *r* = 0.20, *p* < 0.001 but did not differ between view and diminish, *t*(104) = 1.20, *r* = 0.05, *p* = 0.23. Activity in the right amygdala did not significantly differ between intensify and view, *t*(104) = 2.04, *r* = 0.10, *p* = 0.04, nor between view and diminish, *t*(104) = 1.67, *r =*0.08, *p* = 0.10, but differed between intensify and view at an uncorrected *p* threshold (see Table 6 for details).

**Table 6.** Activity Difference in Amygdala ROI between Regulation and View Conditions

| Contrast | M | SE | t | df | p | 95% CI |
| --- | --- | --- | --- | --- | --- | --- |
| Left amygdala |  |  |  |  |  |  |
| Intensify > View | 0.032 | 0.008 | 4.118 | 104 | <0.001 | [0.017, 0.048] |
| View > Diminish | 0.008 | 0.006 | 1.202 | 104 | 0.232 | [-0.005, 0.020] |
| Right amygdala |  |  |  |  |  |  |
| Intensify > View | 0.015 | 0.007 | 2.035 | 104 | 0.044 | [0.0004, 0.029] |
| View > Diminish | 0.010 | 0.006 | 1.665 | 104 | 0.099 | [-0.002, 0.023] |

*Note.* Pairwise comparisons were performed on percent signal change values between conditions.

**Figure 7.**
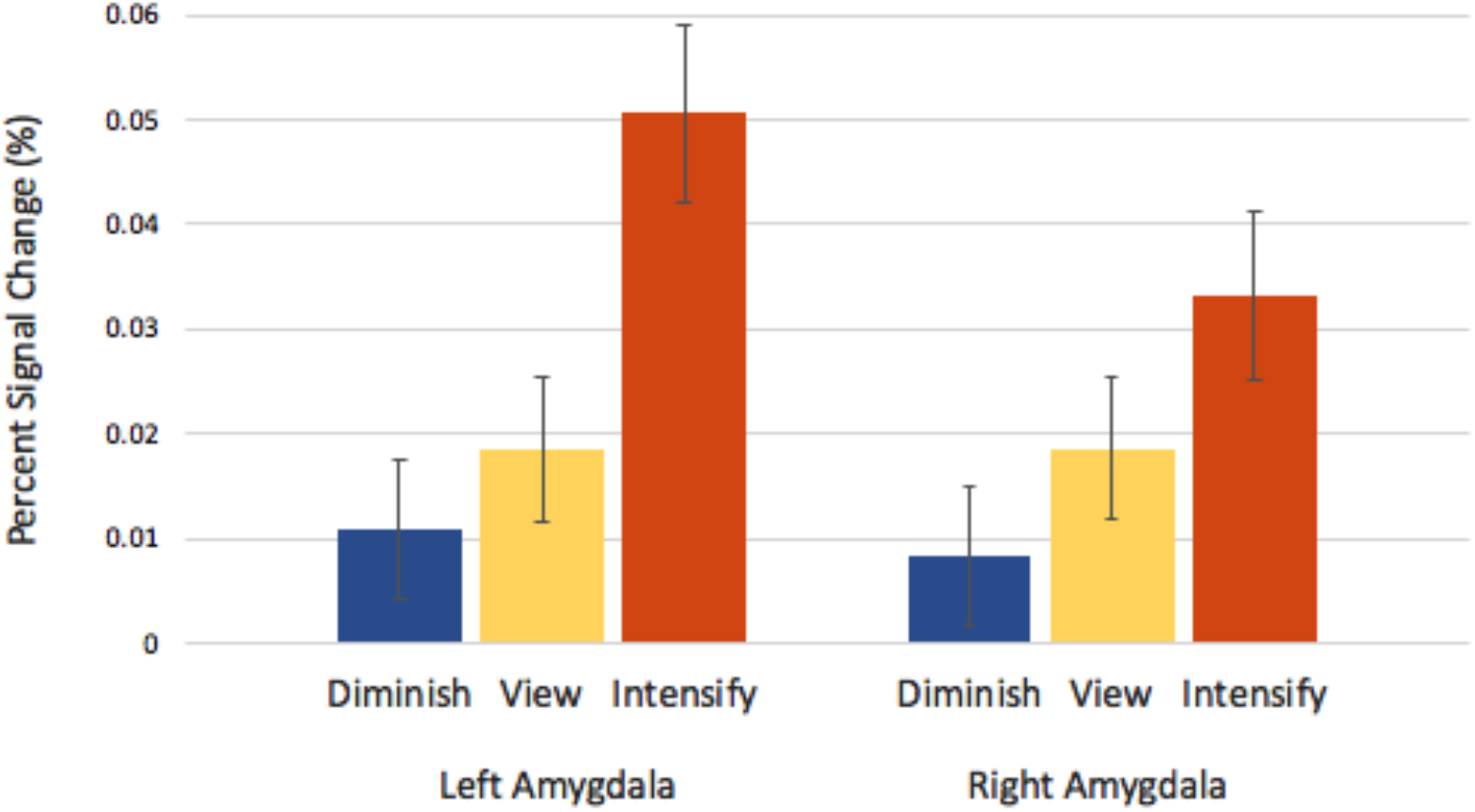
Activity in the amygdala ROIs during down-regulation, viewing and up-regulation *Note.* The error bars reflect the standard error of each condition.

### Regulation Outcome

Our findings that up- and down-regulation effort modulated mostly non-overlapping affect-generating regions (Figure 4B and 4C) raise interesting questions. For instance, which brain regions inform subjective experience of the emotion regulation outcome? And do these also differ during up- and down-regulation? To compare relationships between self-rated intensity and brain activity across up- and down-regulation conditions which differed in average subjective emotion intensity, we normalized rating scores within each of the two regulation conditions for each participant and used these normalized scores as a parametric regressor. Thus, this regressor weighted each trial based on how extreme each participant’s intensity rating was on that trial compared to the average rating for diminishing or intensifying trials. The standard deviation of raw ratings did not significantly differ between intensifying and diminishing trials, *t*(95) = 1.125, *r* = 0.06, *p* = 0.26, indicating similar variability in emotional intensity in the two conditions.

We first examined brain regions whose activity during the 6-second task period (Figure 2) was positively associated with self-rated emotional intensity separately for each condition. While higher subjective ratings after diminishing were associated with the anterior cingulate and paracingulate gyrus (Figure 8A, Table 7), the ratings after intensifying were associated with broader areas including the dorsal anterior cingulate gyrus (ACC), supplementary motor cortex, lingual gyrus, thalamus, and cerebellum (Figure 8B, Table 8). The dorsal ACC, insula, thalamus, and frontal pole were overlapping areas that were associated with greater subjective emotional intensity across both intensifying and diminishing conditions (Figure 9A). We then examined whether there were any brain regions in which activity was negatively associated with self-rated emotional intensity. There were no significant regions for the diminish condition (Figure 8C), but in the intensify condition, there was less activity in right frontoparietal regions during trials with higher self-rated intensity (Figure 8D, Table 9).

**Table 7.** List of Regions (Figure 8A) which Increased Activity as Subjective Ratings Increased during Diminish Trials

| Regions positively correlated with ratings during diminish | MNI coordinate |  |  | Zmax | Voxels |
| --- | --- | --- | --- | --- | --- |
|  | x | y | z |  |  |
| Cingulate Gyrus, anterior division | -2 | 28 | 24 | 4.97 | 618 |
| Insular Cortex | -34 | 12 | 4 | 4.88 | 135 |
| Right Thalamus | 4 | -6 | 0 | 4.50 | 78 |
| Frontal Operculum Cortex | 40 | 24 | 2 | 4.20 | 34 |
| Middle Frontal Gyrus | -30 | 30 | 34 | 4.14 | 50 |
| Paracingulate Gyrus | 6 | 18 | 38 | 4.13 | 82 |
| Supramarginal Gyrus, posterior division | 64 | -40 | 18 | 3.92 | 50 |
| Supplementary Motor Cortex | -4 | 6 | 48 | 3.82 | 11 |
| Frontal Pole | -4 | 58 | 0 | 3.57 | 53 |
| Left Thalamus | 0 | -8 | 6 | 3.49 | 13 |
| Supramarginal Gyrus, anterior division | 64 | -30 | 30 | 3.48 | 2 |
| Central Opercular Cortex | 48 | 6 | 2 | 3.25 | 2 |
| Frontal Medial Cortex | -4 | 54 | -8 | 3.10 | 1 |

**Table 8.**
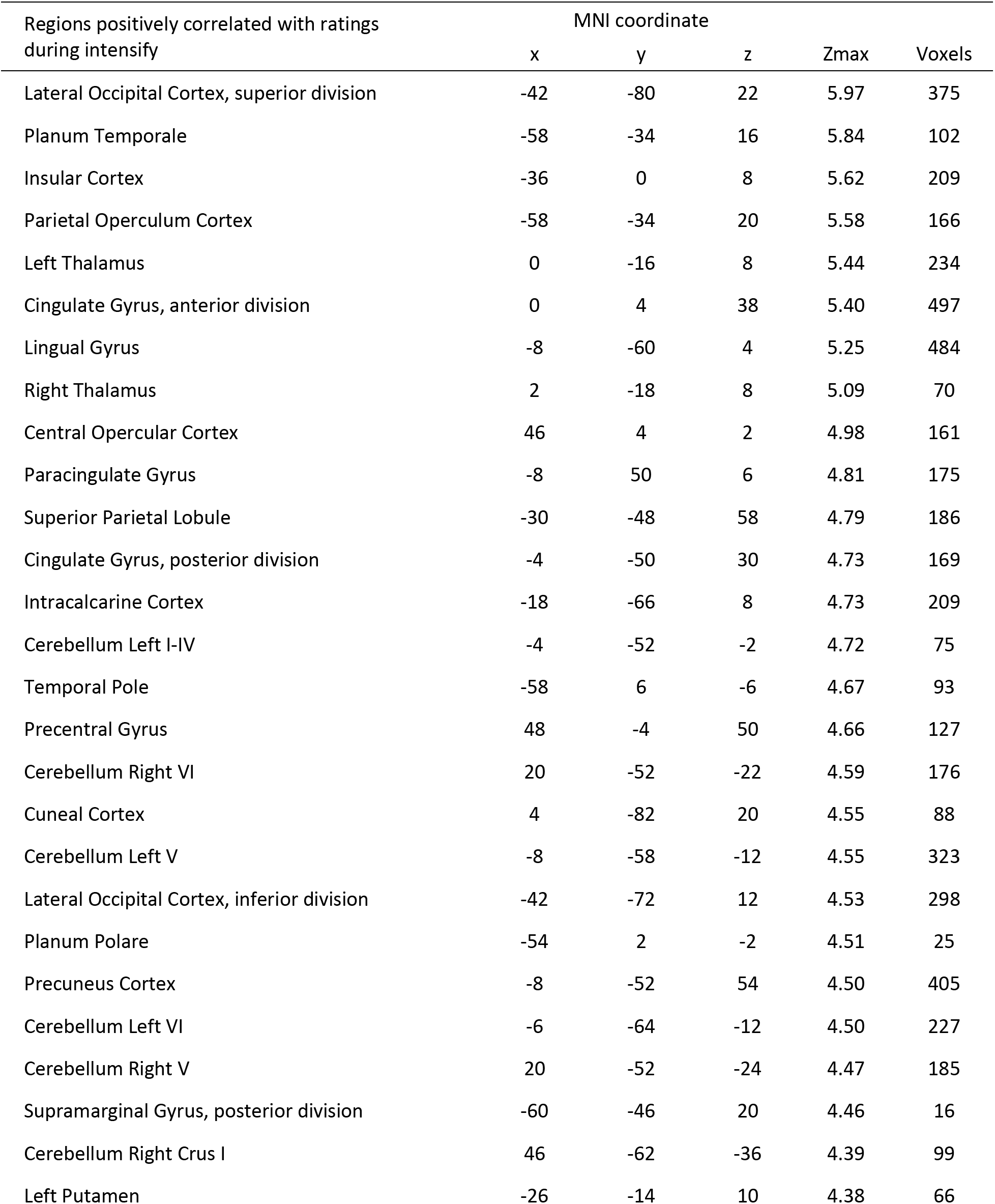
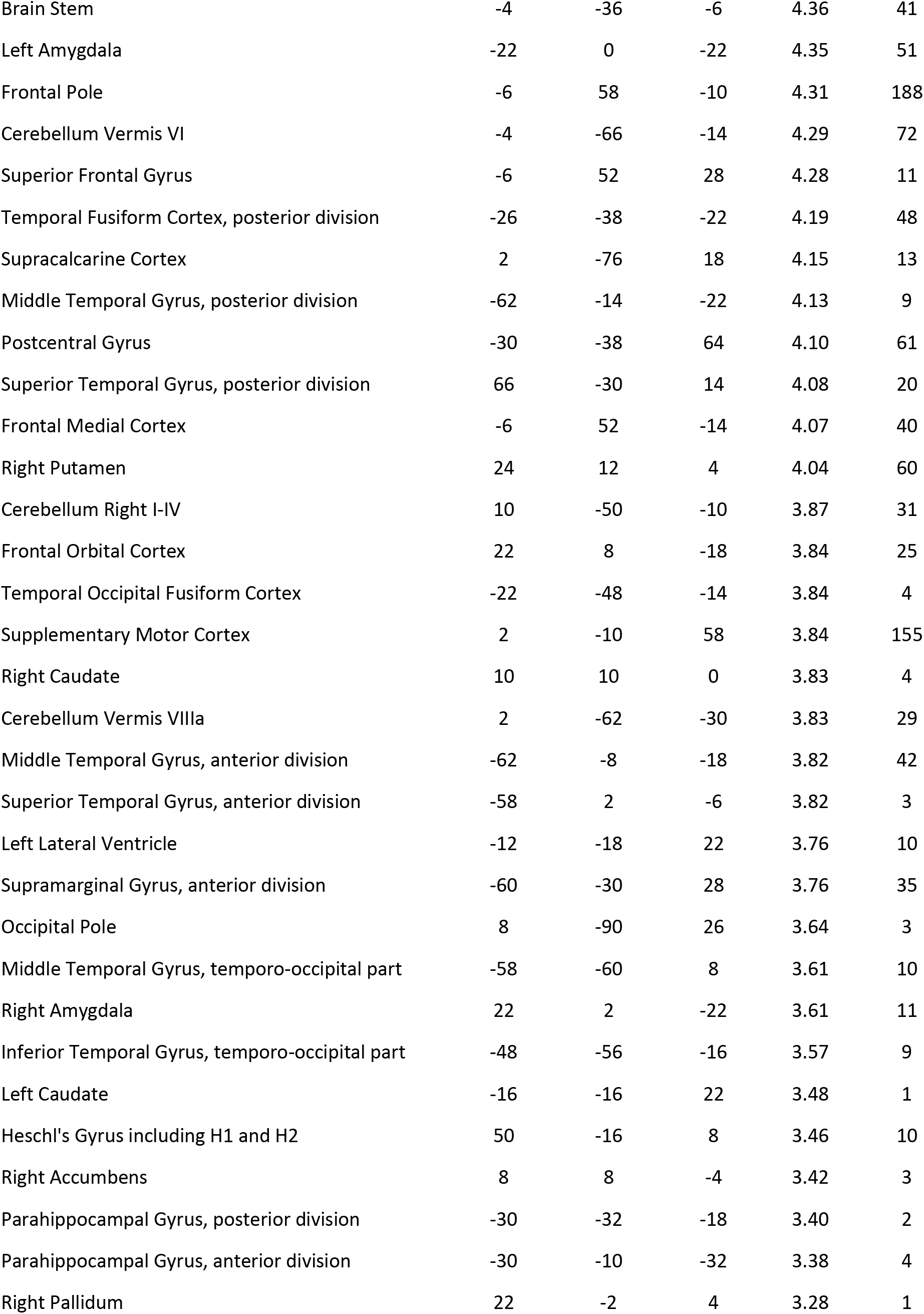
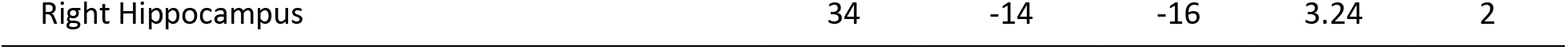
List of Regions (Figure 8B) which Increased Activity as Subjective Ratings Increased during Intensify Trials

**Table 9.** List of Regions (Figure 8D) which Decreased Activity as Subjective Ratings Increased during Intensify Trials

| Regions negatively correlated with ratings during intensify | MNI coordinate |  |  | Zmax | Voxels |
| --- | --- | --- | --- | --- | --- |
|  | x | y | z |  |  |
| Angular Gyrus | 50 | -56 | 42 | 6.43 | 321 |
| Cingulate Gyrus, anterior division | 4 | 28 | 30 | 3.57 | 1 |
| Frontal Pole | 40 | 56 | 2 | 6.11 | 646 |
| Lateral Occipital Cortex, superior division | 44 | -64 | 42 | 6.21 | 487 |
| Middle Frontal Gyrus | 42 | 26 | 38 | 5.72 | 307 |
| Paracingulate Gyrus | 4 | 26 | 44 | 5.14 | 276 |
| Precuneus Cortex | 12 | -68 | 32 | 4.56 | 55 |
| Superior Frontal Gyrus | 20 | 26 | 56 | 4.81 | 40 |
| Supramarginal Gyrus, posterior division | 52 | -44 | 46 | 5.42 | 67 |

**Figure 8.**
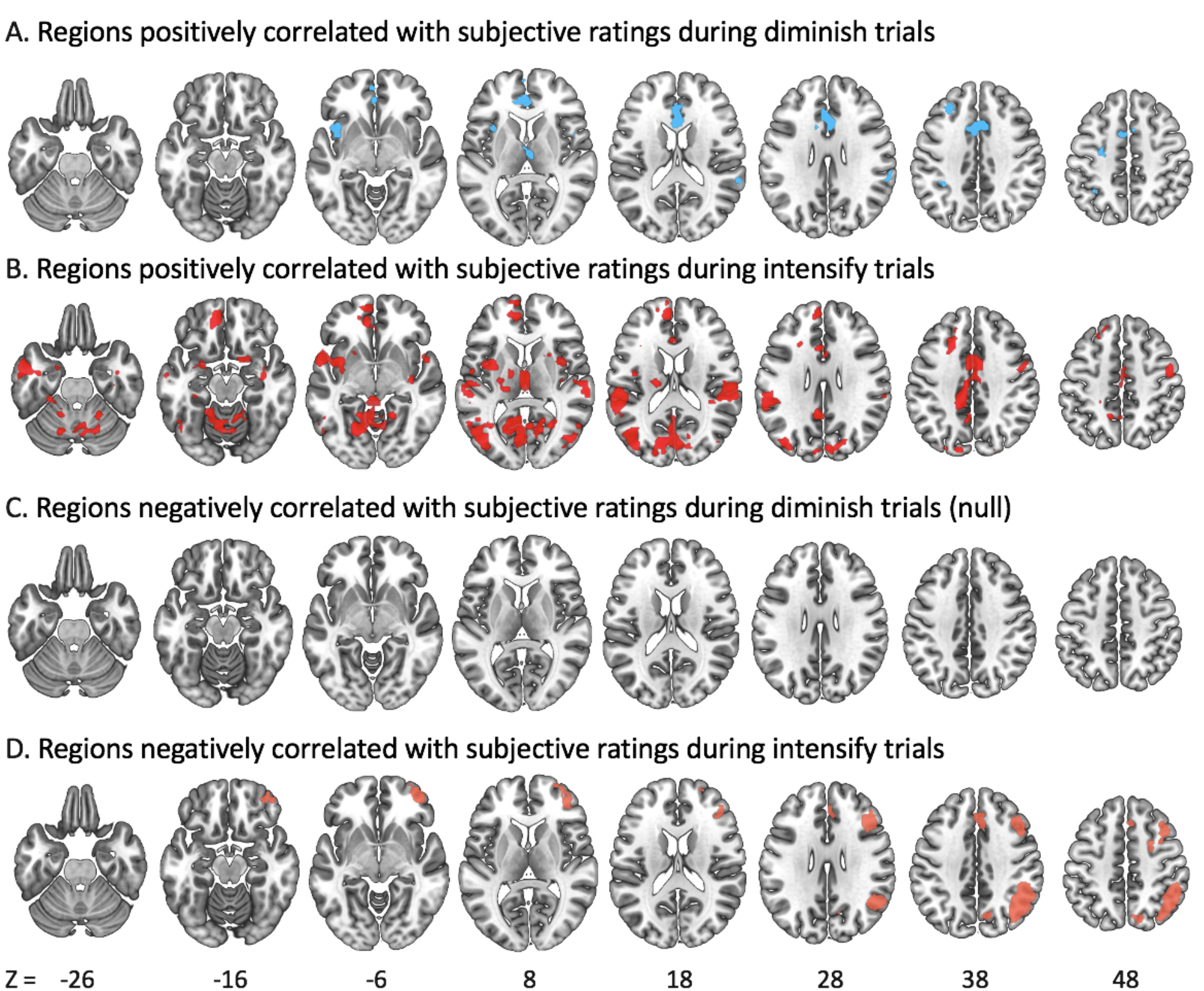
Regions Correlated with Subjective Intensity Ratings during Diminish or Intensify Trials *Note.* (A) shows regions (blue) which increased activity as subjective ratings increased during diminish trials. (B) shows regions (red) which increased activity as subjective ratings increased during intensify trials. (C) would have shown regions (null) which increased activity as subjective ratings decreased during diminish trials. (D) shows regions (orange) which decreased activity as subjective ratings increased during intensify trials.

**Figure 9.**
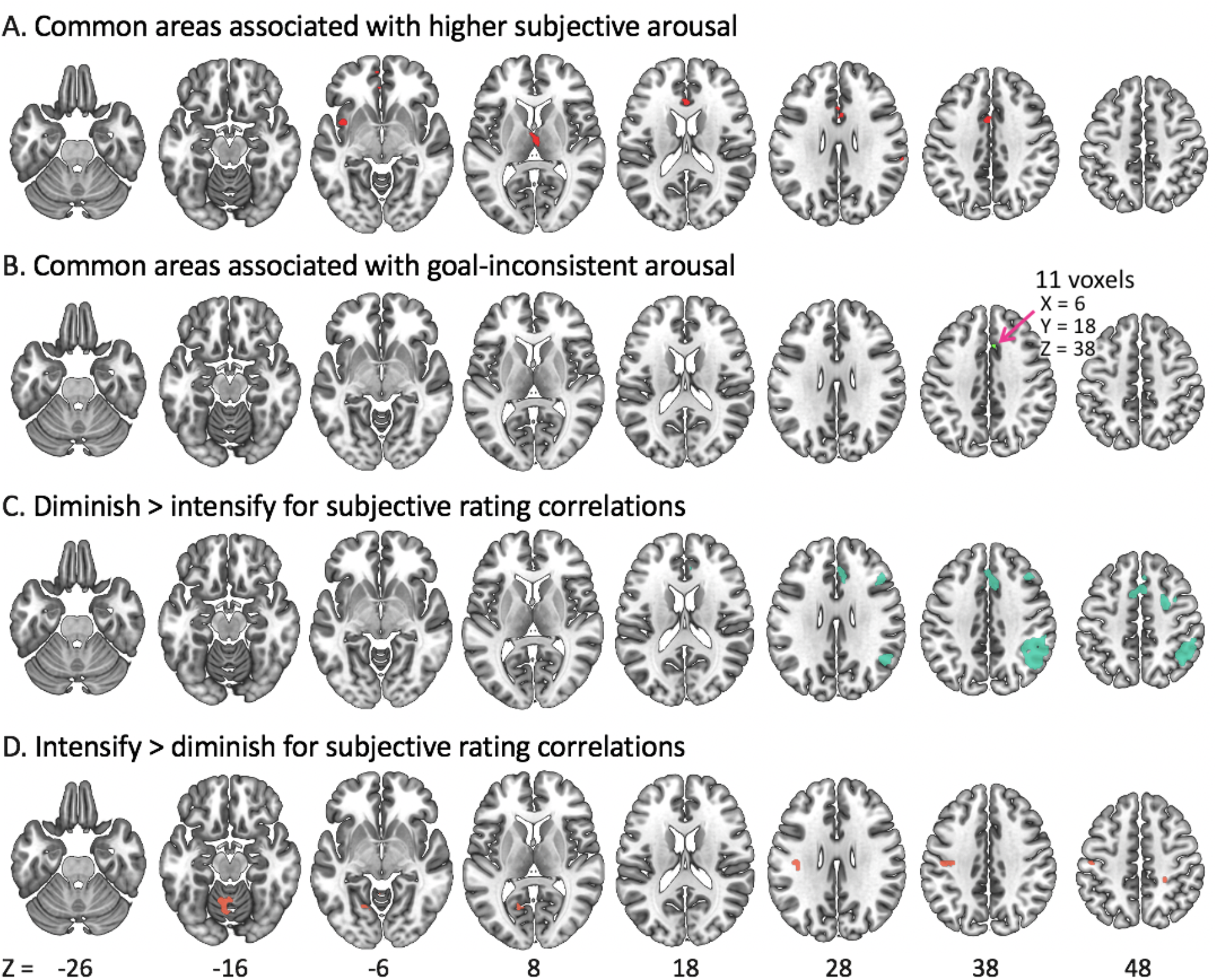
Similarities and Differences Between Regulation Conditions in the Regions Correlated with Subjective Intensity Ratings *Note.* (A) shows the intersection (red) of regions positively correlated with subjective ratings during diminish trials (Figure 8A) and during intensify trials (Figure 8B). (B) shows the intersection (mint) of regions positively correlated with subjective ratings during diminish trials (Figure 8A) and regions negatively correlated with subjective ratings during intensify trials (Figure 8D). (C) shows regions (green) correlated with subjective ratings more positively during diminish than intensify trials or more negatively during intensify than diminish trials. (D) shows regions (orange) correlated with subjective ratings more positively during intensify than diminish trials or more negatively during diminish than intensify trials.

The intersection of Figures 8A and 8B revealed that during both up- and down-regulation, participants reported greater feeling intensity when activation in the insula, ACC, and thalamus were higher (Figure 9A). In contrast, the intersection of Figures 8A and 8D reflects goal-inconsistent arousal in both conditions (i.e., higher feeling intensity during diminish trials and lower feeling intensity during intensify trials) and revealed a separate ACC region (Figure 9B). There also were some significant differences across regulation conditions in how self-perceived emotional intensity was associated with brain activity. The diminish > intensify contrast revealed significant condition differences in the angular gyrus, supramarginal gyrus, dorsal anterior cingulate gyrus, paracingulate gyrus, and middle frontal gyrus (Figure 9C). The intensify > diminish contrast revealed significant differences in the postcentral gyrus and superior parietal lobule (Figure 9D). However, it is important to note that both of the differences across regulation conditions were driven by effects within the intensify condition, as the regions in 9C overlap with those in 8D, which indicates greater negative associations between frontoparietal regions and intensity ratings during intensify than diminish trials, and those in 9D overlap with those in 8B. Thus, we did not find any evidence of regions that are more associated with subjective intensity during diminishing than during intensifying emotions.

## Discussion

The idea that exerting emotional control increases activity in affect-generating brain regions during emotion up-regulation and decreases activity in those same regions during emotion down-regulation makes intuitive sense. This ‘affective dial hypothesis’ is an implicit assumption in the field of emotion regulation. However, our well-powered (N=105) study demonstrated that up- and down-regulation target separate brain regions. The majority of brain regions down-regulated by diminishing did not overlap with those up-regulated by intensifying emotions, as indicated by the minimal intersection between the intensify > view and view > diminish contrasts (Figure 5C).

The intensify > view contrast showed increased activity during up-regulation in many brain regions (Figure 4B) previously associated with affective experience, including the amygdala, anterior insular cortex, ACC, thalamus and nucleus accumbens as well as in regions associated with sympathetic vascular activity such as periventricular white matter (Özbay et al., 2018). Instead of showing decreased activity in these same brain regions during down-regulation as would be predicted by the affective dial hypothesis, the view > diminish contrast revealed decreased activity in the posterior insular cortex and postcentral gyrus (Figure 4C). These areas receive visceral information through the afferent vagus nerve and are involved in interoceptive awareness (Craig, 2002; Khalsa, Rudrauf, Feinstein, & Tranel, 2009). Indeed, more of the voxels activated during down-regulation overlapped brain regions that a previous meta-analysis (Adolfi et al., 2017) linked with interoception than overlapped brain regions linked with other aspects of emotion whereas the reverse was the case for up-regulation (Figure 6D). Why might up- and down-regulation target different affective circuits? When up-regulating, people may engage more with emotional images, whereas when down-regulating they may disengage more. Engaging with external emotional stimuli may target emotional processing pathways that help evaluate external stimuli, while disengaging from external stimuli with the goal of reducing feelings may instead target interoceptive processing pathways. Future research should test these and other possibilities.

Even though the brain regions targeted by up- and down-regulation barely overlapped, these regulatory modes activated an overlapping set of brain regions (that is, overlap in intensify > view and diminish > view; Figure 5A). These overlapping regions included the inferior frontal gyrus, dorsal anterior cingulate gyrus (ACC), and anterior insular cortex, which have been previously linked with various aspects of emotion regulation. Our participants used cognitive reappraisal strategies; the inferior frontal gyrus (IFG), within the ventrolateral prefrontal cortex (vlPFC), is involved in strategies which require modifying interpretations of emotional situations to attenuate negative emotion (Ochsner & Gross, 2005). The dorsal ACC detects conflicts and signals adjustments in cognitive tasks (Botvinick, Cohen, & Carter, 2004; Bush, Luu, & Posner, 2000) and emotion regulation (Etkin, Büchel, & Gross, 2015; Ichikawa et al., 2011; McRae, Reiman, Fort, Chen, & Lane, 2008). Anterior insula activation is associated with subjective feelings of emotion and their autonomic representation (Craig, 2009; Critchley & Harrison, 2013).

The amygdala showed a linear pattern where its BOLD activity was highest during up-regulation, mid-range during viewing, and lowest during down-regulation (Figure 7). This seemed to support prior work which has focused on how emotion regulation modulates amygdala activity (e.g., Goldin, McRae, Ramel, & Gross, 2008; Kim & Hamann, 2007; McRae et al., 2010; Ochsner et al., 2004; Steinfurth et al., 2018). However, the amygdala did not show any significant voxels in the affective-dial intensify > view ∩ view > diminish contrast at the whole-brain level with our conservative threshold (cluster size *Z* > 3.1). To our knowledge, there are no prior findings of overlapping up-regulation > baseline and baseline > down-regulation effects in the amygdala at a whole-brain threshold level. Prior studies typically employed region-of-interest (ROI) or small-volume-corrected analyses, allowing for lenient statistical thresholds. They also relied on relatively small numbers of participants (e.g., N = 10 – 24).

Further examination of our whole-brain results suggests that our conjunction analysis did not reveal an affective-dial pattern in the amygdala because the view > diminish and intensify > view contrasts activated different parts of the amygdala. The view > diminish contrast activated its laterobasal subregions, whereas the intensify > view contrast activated mostly the superficial and centromedial subregions. While the laterobasal subregion of the amygdala receives sensory information from the visual and auditory cortex, the centromedial subregion is related to emotional arousal and responses (Kerestes, Chase, Phillips, Ladouceur, & Eickhoff, 2017). Future research should investigate whether up- and down-regulation do indeed target different amygdala subregions.

During the last 4 seconds of each trial, participants rated the intensity of feelings (Figure 2). These ratings were lower in the diminish than in the intensify condition (Figure 3). But do lower vs. higher ratings relate to activity in the same brain regions during up- versus down- regulating emotion? Indeed, we found several brain regions where increased activity both during diminishing and intensifying emotions were significantly associated with relatively greater intensity ratings (Figure 9A). These included the left insula (Figure 9A) and a small cluster in the right insula (not shown). The insula’s activity level may help signal affective intensity as it is associated with both interoception and other aspects of emotion (Figure 6C, Adolfi et al., 2017). Other regions where activity was associated with subjective intensity included the dorsal ACC and the frontal pole, which, as part of the medial PFC, activate during self-referencing tasks involving emotional stimuli (Northoff et al., 2006).

There were also some interesting differences across conditions. When the goal was to intensify emotions, higher subjective feeling intensity was associated with lower activity in the right frontoparietal attention network (e.g., Laird et al., 2011) during up-regulation, suggesting that intensifying emotions suppresses activity in this attention network (Figure 8D). Directly contrasting the correlations with subjective feelings in the two conditions revealed that this suppression of frontoparietal activity was significantly more associated with subjective feelings during intensifying than during diminishing emotion (Figure 9C, 9D). Thus, whereas amping up emotion during up-regulation suppresses frontoparietal activity (Figure 8D), tamping down emotion during down-regulation does not increase frontoparietal activity (Figure 8C).

In contrast, activity in a dorsal ACC region (Figure 9B) during emotion regulation was associated with lower intensity ratings during intensify trials (that is, a failure to achieve the instructed higher arousal state; Figure 8D) and with higher intensity ratings on diminish trials (that is, again, a failure to achieve the instructed lower arousal state; Figure 8A). This region appears to be providing a task-failure signal (or reflecting compensatory effort in response to failure), consistent with the role of the dorsal ACC in error monitoring (Gilbertson, Fang, Andrzejewski, & Carlson, 2021; Taylor, Stern, & Gehring, 2007). Thus, up- and down-regulation appear to rely on some overlapping brain regions (Figure 9B) to integrate arousal signals and to monitor the gap between the goal and actual states, despite the differences identified earlier in affect-generating brain regions targeted by these two regulatory goals.

We observed broad activation in the white matter surrounding the ventricles during intensifying emotion compared to viewing emotional images (Figure 4B). Although we could not find prior studies which explicitly discussed white matter activation during emotion regulation, we observed it in the figures of studies reporting on emotion up-regulation and the viewing of highly emotional images (e.g., Grosse Rueschkamp et al., 2019, Figure 4; Moodie et al., 2020, Figure 3). Increased white matter BOLD signal associated with increased emotional arousal might be due to sympathetic activity increasing vascular tone (Özbay et al., 2018). White matter veins converge to subependymal veins that run around the edge of the lateral ventricles (Okudera et al., 1999), and so periventricular white matter is especially susceptible to systemic changes in vascular tone (Özbay et al., 2018). Blood oxygen level dependent (BOLD) signal is weaker in white matter than in grey matter (Gawryluk, Mazerolle, & D’Arcy, 2014), but our use of a multi-echo sequence to remove noise components and our large N likely provided stronger power than prior studies to detect such effects. The vascular aspect of BOLD signals associated with emotional arousal has yet to be fully explored in the field of emotion regulation. Future studies should examine how the autonomic nervous system interacts with vascular mechanisms and how that interaction affects brain activity during emotion regulation.

In summary, the current study investigated brain regions associated with emotion regulation by employing cognitive reappraisal strategies and demonstrated that up- and down-regulation exert control on distinct brain regions. The regions targeted by up-regulation were more likely to be involved in emotional arousal whereas regions targeted by down-regulation were more likely to be involved in interoception. These findings indicate that up- and down-regulating our emotions using cognitive reappraisal are not simply mirror image processes that have opposing effects on the same emotion-generating brain regions. Instead, they target different affective circuits in the brain. As such, our findings raise the possibility that some individuals may excel at up- but not at down-regulating their own emotions, or vice versa.

## Acknowledgements

This study was supported by NIH R01AG057184 (PI Mather). We thank our research assistants for their help with data collection: Linette Bagtas, Akanksha Jain, Divya Suri, Sophia Ling, Michelle Wong, Yong Zhang and Gabriel Shih.

## References

Adolfi, F., Couto, B., Richter, F., Decety, J., Lopez, J., Sigman, M., Manes, F., & Ibáñez, A. (2017). Convergence of interoception, emotion, and social cognition: a twofold fMRI meta-analysis and lesion approach. Cortex, 88, 124–142.

Berboth, S., & Morawetz, C. (2021). Amygdala-prefrontal connectivity during emotion regulation: A meta-analysis of psychophysiological interactions. Neuropsychologia, 153, 107767.

Botvinick, M. M., Cohen, J. D., & Carter, C. S. (2004). Conflict monitoring and anterior cingulate cortex: an update. Trends in Cognitive Sciences, 8(12), 539–546.

Braunstein, L. M., Gross, J. J., & Ochsner, K. N. (2017). Explicit and implicit emotion regulation: a multi-level framework. Social cognitive and affective neuroscience, 12(10), 1545–1557.

Buhle, J. T., Silvers, J. A., Wager, T. D., Lopez, R., Onyemekwu, C., Kober, H., Weber, J., & Ochsner, K. N. (2014). Cognitive reappraisal of emotion: a meta-analysis of human neuroimaging studies. Cerebral cortex, 24(11), 2981–2990.

Bush, G., Luu, P., & Posner, M. I. (2000). Cognitive and emotional influences in anterior cingulate cortex. Trends in Cognitive Sciences, 4(6), 215–222.

Craig, A. D. (2002). How do you feel? Interoception: the sense of the physiological condition of the body. Nature reviews neuroscience, 3(8), 655–666.

Craig, A. D. (2009). How do you feel--now? The anterior insula and human awareness. Nature reviews neuroscience, 10(1).

Critchley, H. D., & Harrison, N. A. (2013). Visceral influences on brain and behavior. Neuron, 77(4), 624–638.

Domes, G., Schulze, L., Böttger, M., Grossmann, A., Hauenstein, K., Wirtz, P. H., Heinrichs, M., & Herpertz, S. C. (2010). The neural correlates of sex differences in emotional reactivity and emotion regulation. Human brain mapping, 31(5), 758–769.

Eippert, F., Veit, R., Weiskopf, N., Erb, M., Birbaumer, N., & Anders, S. (2007). Regulation of emotional responses elicited by threat-related stimuli. Human brain mapping, 28(5), 409–423.

Etkin, A., Büchel, C., & Gross, J. J. (2015). The neural bases of emotion regulation. Nature reviews neuroscience, 16(11), 693–700.

Gawryluk, J. R., Mazerolle, E. L., & D’Arcy, R. C. (2014). Does functional MRI detect activation in white matter? A review of emerging evidence, issues, and future directions. Frontiers in neuroscience, 8, 239.

Gilbertson, H., Fang, L., Andrzejewski, J. A., & Carlson, J. M. (2021). Dorsal anterior cingulate cortex intrinsic functional connectivity linked to electrocortical measures of error monitoring. Psychophysiology, 58(5), e13794.

Goldin, P. R., McRae, K., Ramel, W., & Gross, J. J. (2008). The neural bases of emotion regulation: reappraisal and suppression of negative emotion. Biological psychiatry, 63(6), 577–586.

Gross, J. J. (2015). Emotion regulation: Current status and future prospects. Psychological inquiry, 26(1), 1–26.

Grosse Rueschkamp, J. M., Brose, A., Villringer, A., & Gaebler, M. (2019). Neural correlates of up-regulating positive emotions in fMRI and their link to affect in daily life. Social cognitive and affective neuroscience, 14(10), 1049–1059.

Ichikawa, N., Siegle, G. J., Jones, N. P., Kamishima, K., Thompson, W. K., Gross, J. J., & Ohira, H. (2011). Feeling bad about screwing up: emotion regulation and action monitoring in the anterior cingulate cortex. Cognitive, Affective, & Behavioral Neuroscience, 11(3), 354–371.

Kerestes, R., Chase, H. W., Phillips, M. L., Ladouceur, C. D., & Eickhoff, S. B. (2017). Multimodal evaluation of the amygdala’s functional connectivity. Neuroimage, 148, 219–229.

Khalsa, S. S., Rudrauf, D., Feinstein, J. S., & Tranel, D. (2009). The pathways of interoceptive awareness. Nature neuroscience, 12(12), 1494–1496.

Kim, S. H., & Hamann, S. (2007). Neural correlates of positive and negative emotion regulation. Journal of cognitive neuroscience, 19(5), 776–798.

Kohn, N., Eickhoff, S. B., Scheller, M., Laird, A. R., Fox, P. T., & Habel, U. (2014). Neural network of cognitive emotion regulation—an ALE meta-analysis and MACM analysis. Neuroimage, 87, 345–355.

Kundu, P., Inati, S. J., Evans, J. W., Luh, W.-M., & Bandettini, P. A. (2012). Differentiating BOLD and non-BOLD signals in fMRI time series using multi-echo EPI. Neuroimage, 60(3), 1759–1770.

Laird, A. R., Fox, P. M., Eickhoff, S. B., Turner, J. A., Ray, K. L., McKay, D. R., Glahn, D. C., Beckmann, C. F., Smith, S. M., & Fox, P. T. (2011). Behavioral interpretations of intrinsic connectivity networks. Journal of cognitive neuroscience, 23(12), 4022–4037.

Leiberg, S., Eippert, F., Veit, R., & Anders, S. (2012). Intentional social distance regulation alters affective responses towards victims of violence: an FMRI study. Human brain mapping, 33(10), 2464–2476.

Li, F., Yin, S., Feng, P., Hu, N., Ding, C., & Chen, A. (2018). The cognitive up-and down-regulation of positive emotion: Evidence from behavior, electrophysiology, and neuroimaging. Biological psychology, 136, 57–66.

Lindquist, K. A., Satpute, A. B., Wager, T. D., Weber, J., & Barrett, L. F. (2016). The brain basis of positive and negative affect: evidence from a meta-analysis of the human neuroimaging literature. Cerebral cortex, 26(5), 1910–1922.

McRae, K., Hughes, B., Chopra, S., Gabrieli, J. D., Gross, J. J., & Ochsner, K. N. (2010). The neural bases of distraction and reappraisal. Journal of cognitive neuroscience, 22(2), 248–262.

McRae, K., Reiman, E. M., Fort, C. L., Chen, K., & Lane, R. D. (2008). Association between trait emotional awareness and dorsal anterior cingulate activity during emotion is arousal-dependent. Neuroimage, 41(2), 648–655.

Moodie, C. A., Suri, G., Goerlitz, D. S., Mateen, M. A., Sheppes, G., McRae, K., Lakhan-Pal, S., Thiruchselvam, R., & Gross, J. J. (2020). The neural bases of cognitive emotion regulation: The roles of strategy and intensity. Cognitive, Affective, & Behavioral Neuroscience, 20(2), 387–407.

Morawetz, C., Alexandrowicz, R. W., & Heekeren, H. R. (2017). Successful emotion regulation is predicted by amygdala activity and aspects of personality: A latent variable approach. Emotion, 17(3), 421.

Morawetz, C., Bode, S., Baudewig, J., Jacobs, A. M., & Heekeren, H. R. (2016). Neural representation of emotion regulation goals. Human brain mapping, 37(2), 600–620.

Morawetz, C., Bode, S., Baudewig, J., Kirilina, E., & Heekeren, H. R. (2016). Changes in effective connectivity between dorsal and ventral prefrontal regions moderate emotion regulation. Cerebral cortex, 26(5), 1923–1937.

Morawetz, C., Bode, S., Derntl, B., & Heekeren, H. R. (2017). The effect of strategies, goals and stimulus material on the neural mechanisms of emotion regulation: a meta-analysis of fMRI studies. Neuroscience & Biobehavioral Reviews, 72, 111–128.

Northoff, G., Heinzel, A., De Greck, M., Bermpohl, F., Dobrowolny, H., & Panksepp, J. (2006). Self-referential processing in our brain—a meta-analysis of imaging studies on the self. Neuroimage, 31(1), 440–457.

Ochsner, K. N., & Gross, J. J. (2005). The cognitive control of emotion. Trends in Cognitive Sciences, 9(5), 242–249.

Ochsner, K. N., Ray, R. D., Cooper, J. C., Robertson, E. R., Chopra, S., Gabrieli, J. D., & Gross, J. J. (2004). For better or for worse: neural systems supporting the cognitive down-and up-regulation of negative emotion. Neuroimage, 23(2), 483–499.

Ochsner, K. N., Silvers, J. A., & Buhle, J. T. (2012). Functional imaging studies of emotion regulation: a synthetic review and evolving model of the cognitive control of emotion. Annals of the new York Academy of Sciences, 1251, E1.

Okudera, T., Huang, Y. P., Fukusumi, A., Nakamura, Y., Hatazawa, J., & Uemura, K. (1999). Micro-angiographical studies of the medullary venous system of the cerebral hemisphere. Neuropathology, 19(1), 93–111.

Özbay, P. S., Chang, C., Picchioni, D., Mandelkow, H., Chappel-Farley, M. G., van Gelderen, P., de Zwart, J. A., & Duyn, J. (2019). Sympathetic activity contributes to the fMRI signal. Communications biology, 2(1), 1–9.

Özbay, P. S., Chang, C., Picchioni, D., Mandelkow, H., Moehlman, T. M., Chappel-Farley, M. G., van Gelderen, P., de Zwart, J. A., & Duyn, J. H. (2018). Contribution of systemic vascular effects to fMRI activity in white matter. Neuroimage, 176, 541–549.

Phelps, E. A. (2006). Emotion and cognition: insights from studies of the human amygdala. Annu. Rev. Psychol., 57, 27–53.

Steinfurth, E. C., Wendt, J., Geisler, F., Hamm, A. O., Thayer, J. F., & Koenig, J. (2018). Resting state vagally-mediated heart rate variability is associated with neural activity during explicit emotion regulation. Frontiers in neuroscience, 12, 794.

Taylor, S. F., Stern, E. R., & Gehring, W. J. (2007). Neural systems for error monitoring: recent findings and theoretical perspectives. The Neuroscientist, 13(2), 160–172.

